# Higher-Order Organization Principles of Pre-translational mRNPs

**DOI:** 10.1101/278747

**Authors:** Mihir Metkar, Hakan Ozadam, Bryan R. Lajoie, Maxim Imakaev, Leonid A. Mirny, Job Dekker, Melissa J. Moore

**Author notes:** Current Affiliation: Illumina, San Diego, CA. Current Affiliation: Moderna Therapeutics, Cambridge, MA.

## Abstract

Compared to noncoding RNAs (ncRNAs) such as rRNAs and ribozymes, for which high resolution structures abound, little is known about the tertiary structures of mRNAs. In eukaryotic cells, newly made mRNAs are packaged with proteins in highly compacted mRNPs, but the manner of this mRNA compaction is unknown. Here we developed and implemented RIPPLiT (<u>R</u>NA Immuno<u>P</u>recipitation and Proximity Ligation in Tandem), a transcriptome-wide method for probing the 3D conformations of RNAs stably-associated with defined proteins, in this case exon junction complex (EJC) core factors. EJCs multimerize with other mRNP components to form megadalton sized complexes that protect large swaths of newly synthesized mRNAs from endonuclease digestion. Unlike ncRNAs, mRNAs behave more like flexible polymers without strong locus-specific interactions. Polymer analysis of proximity ligation data for hundreds of mRNA species demonstrates that pre-translational mammalian mRNPs fold as linear rod-like structures with no strong propensity for 5’ and 3’ end interaction.

## Introduction

Once synthesized, messenger RNA particles (mRNPs) must explore the nuclear and cytoplasmic compartments to arrive at their final subcellular destinations. Such travel necessitates packaging in a manner that prevents RNA tangling and shearing as well as premature degradation. To date, however, the rules governing RNA polymerase II (Pol II) transcript packaging remain largely undefined. Assays such as icSHAPE and DMS-Seq that detect nucleotide accessibility suggest that mRNAs are generally flexible and unstructured, even more so *in vivo* than *in vitro* (Rouskin et al., 2014; Spitale et al., 2015). Such studies, however, provide no information regarding the conformational properties of the RNA polymer inside the mRNP or its degree of compaction. Conformationally, the RNA could fold as a simple random coil, an equilibrium globule, a fractal globule, or some other polymer arrangement, and this unstructured arrangement could be either loosely packed or highly compacted.

The only available data about overall mRNP shape in cells come from electron microscopy (EM) studies where images of purified polyA+ transcripts from budding yeast (in which mRNAs average ~1250 nt (Miura et al., 2008)) revealed rod-like structures of differing lengths but nearly constant width (~5 nm) (Batisse et al., 2009). At the opposite extreme, *in situ* EM images of giant Balbiani ring mRNAs (35,000 to 40,000 nt) in *Chironomus tentans* salivary gland nuclei also showed rod-like structures (~10 nm wide) that collapse into 19 nm stalks during transcription and then 50 nm globular structures upon chromatin release (Skoglund et al., 1983). For both yeast and Balbiani ring mRNPs, these measured dimensions necessitate substantial RNA condensation relative to a simple linear structure, with the estimated compaction being ~11-fold for yeast mRNAs and ~200-fold for Balbiani ring mRNAs. But how this is accomplished and what general principles guide mRNP 3D organization are currently unknown. Further, it is unknown whether the globular structures adopted by Balbiani ring mRNPs are unique to these exceptional long transcripts. That is, are mammalian mRNPs (containing >3,900 nt mRNAs on average; NCBI *Homo sapiens* Annotation Release 108) more rod-like or more globular?

Pre-translational mRNPs contain tightly-bound proteins that accompany the mRNA from the nucleus to the cytoplasm. Chief among these are exon junction complexes (EJCs), composed of three core factors: eIF4AIII, Magoh and Y14. These proteins are deposited on Pol II transcripts upstream of exon junctions during pre mRNA splicing and remain in place until their removal by the first round of translation (Dostie and Dreyfuss, 2002; Lejeune et al., 2002). The mechanism of their sequence-independent deposition essentially locks EJCs in place on the RNA until they are unlocked by a factor associated with elongating ribosomes (Gehring et al., 2009). We previously showed that endogenous EJCs interact both with one another and with other tightly bound mRNP components (e.g., peripheral EJC proteins, serine/arginine (SR)- rich proteins and SR-like proteins) through short and long-range interactions to form RNase-resistant, megadalton-sized complexes containing 30-150 nt protected fragments of spliced mRNAs (Singh et al., 2012). Incredibly, even in the absence of any crosslinking agent, these higher order structures are stable to stringent double IPs and nuclease treatments designed for footprinting assays, and have molecular weights exceeding 2 MDa (for comparison, the large ribosomal subunit has a molecular weight of 1.7 MDa and contains ∼5,000 nt). Thus, EJCs and their associated proteins form a large and stable structural core that packages and protects newly made mRNAs. Here we used RNA Proximity Ligation (Kudla et al., 2011; Ramani et al., 2015) to investigate the higher order structure, polymer compaction and RNA folding principles within this core.

## Results

### RIPPLiT: A method to capture higher order RNA structure in RNPs

We previously developed RNA Protein Immunoprecipation in Tandem (RIPiT) (Singh et al., 2012; Singh et al., 2014) to enable purification of RNP complexes containing specified protein pairs. EJC-containing RNPs can be selectively enriched by first affinity-enriching for one EJC core protein (e.g., FLAG-tagged Magoh), and then immunopurifing a second (e.g., eIF4AIII). Inclusion of a nuclease digestion step between the two affinity steps enables the production of footprints. Here we modified this protocol to include a proximity ligation step after nuclease digestion (**Figure 1A**). Briefly, we immunoprecitated FLAG-tagged EJC-containing particles and fragmented the bound RNA via limited RNase T1 digestion to generate a fragment distribution ranging from 30 to >500 nt (**Figure 1B**). Following conversion of RNA ends to 5′-P and 3′-OH groups (necessary for ligation) during the second immunoaffinity step (**Figure 1A**), T4 RNA Ligase I (Rnl I) addition produced a distinct shift toward larger sized RNA fragments (**Figure 1B**). Protection from phosphatase removal of 5′- ^32^P labels incorporated during the phosphorylation step confirmed that these longer fragments were bona fide ligation products (**Figure S1)**. We call this new method for mapping higher ordered structures within double affinity-purified RNPs RNA Protein Immunoprecipation and Proximity Ligation in Tandem (RIPPLiT).

**Figure 1.**
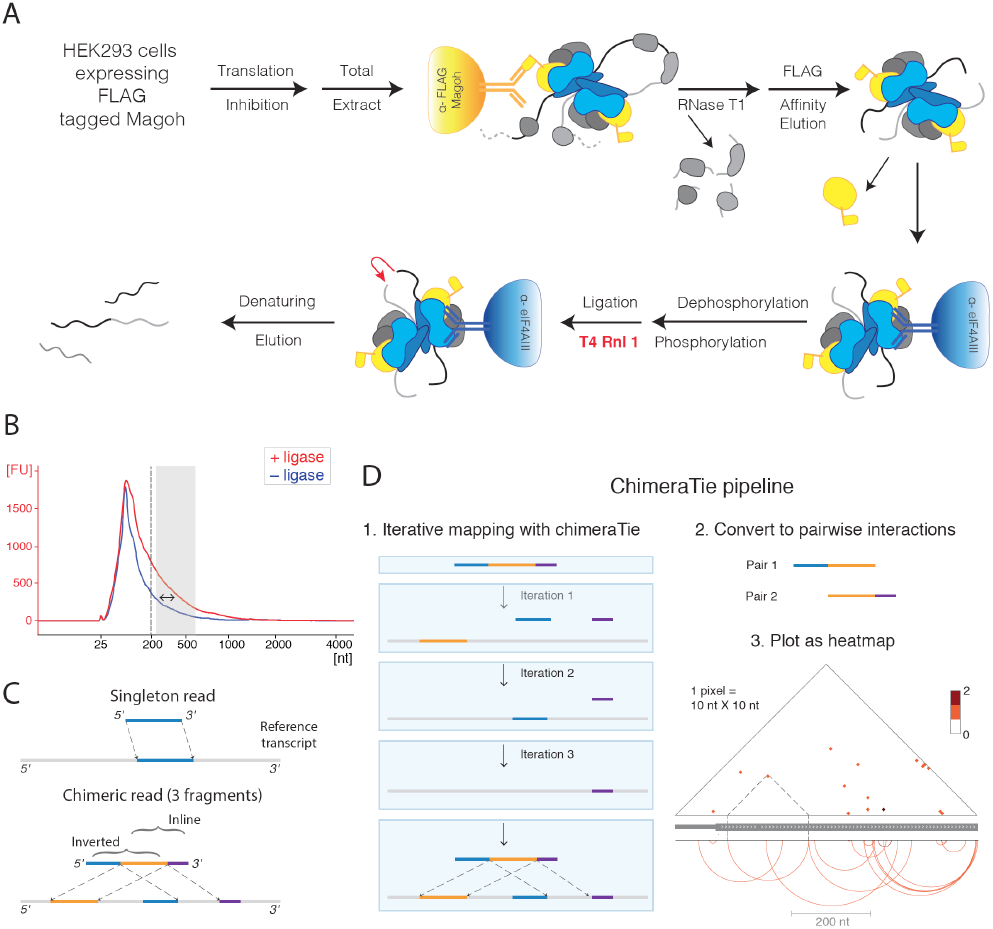
Overview of RIPPLiT and ChimeraTie. (A) Schematic for EJC RIPPLiT. Solid yellow and blue objects, EJC core proteins; gradient yellow and blue objects, antibody-conjugated beads; gray objects, non-EJC proteins; black and grey lines, RNA; red arrow, proximity ligation event. (B) Bioanalyzer trace showing length distribution of RNAs obtained from EJC RIPPLiT (Replicate 1). Double-head arrow, length distribution shift due to addition of ligase; gray box, sizes selected for sequencing. (C) Types of reads, fragments and chimeric junctions in RIPPLiT libraries, and their relationships to reference transcripts. (D) Schematic of ChimeraTie pipeline used to iteratively map fragments, extract pairwise chimeric junctions and then visualize junctions as a heatmap. Data shown are chimeric junctions (Replicate 1 only) within the first 767 nts of PRPF8 mRNA. Thick gray line with arrowheads, coding exons; thinner section; 5’UTR. Arcs show individual chimeric junctions at nt resolution. Heatmap indicates number of junctions within each 10 ×10 nt pixel, with dotted lines indicating the heatmap position of one chimeric junction.

For RNP structural analysis, we performed RIPPLiT on three independent biological replicate whole cell lysates from HEK293 cells expressing FLAG-tagged Magoh. Immediately before the ligation step, each replicate was divided in half, with one half receiving ligase (+ ligase) and the other not (- ligase). All libraries were size selected for ~200-550 nt inserts. Preliminary data analysis by Sanger sequencing revealed that the + ligase libraries contained a high fraction of chimeric reads (i.e., reads made up of concatenated fragments mapping to different locations on one or more RNA species) (**Figure 1C**), whereas the - ligase libraries did not. Paired-end sequencing (150 nt) on the Illumina NextSeq platform yielded 23 to 49 million merged reads per library (**Table S1**). To facilitate chimeric read mapping we developed ChimeraTie, a bioinformatics tool that employs the local alignment mode of Bowtie 2 to iteratively map all fragments within a single read (**Figure 1D**). Pair-wise chimeric junctions are visualized as two-dimensional heatmaps, where color intensities correspond to junction frequencies. Aggregating counts into appropriate length bins assists in data visualization across long RNAs (**Figure 1D.3**).

Confirming our purification of EJC-associated RNAs, 73-79% of fragments in all libraries mapped uniquely to Pol II transcripts (**Table S2A**), with spliced transcripts being significantly enriched over Pol II transcripts containing no annotated intron (**Figure 2A;** 1.05e^−79^, Wilcoxon rank sum test). Whereas both + and - ligase libraries contained similar fractions of individual fragments mapping to Pol II transcripts, intramolecular chimeric junctions were 3.5 to 8-fold higher in + than - libraries (**Figure 2B, inset Table; Table S2A**). In the + ligase libraries, both inline and inverted chimeric junctions were well represented, with both distributed across a wide range of intervening nucleotide distances (aka, spans) (**Figure 2B, cumulative histogram**). In contrast, - ligase chimeric junctions were predominantly inline (**Figure 2B, inset Table**) and these inline junctions were strongly skewed toward shorter spans (**Figure 2B, cumulative histogram**). These features are characteristic of mapping artifacts wherein a long single fragment is mistakenly mapped as two or more shorter inline fragments separated by short gaps. To minimize this issue, we optimized Bowtie2’s read and reference gap penalties to disfavor short fragment junction gaps, and we used only uniquely mapping fragments (i.e., having no secondary alignment) with fewer than 3 mismatches/indels for all analyses. Nonetheless, since the relative weighting of a gap penalty decreases as alignment length increases in Bowtie2, we were unable to completely eliminate all false positive, inline chimeric junctions.

**Figure 2.**
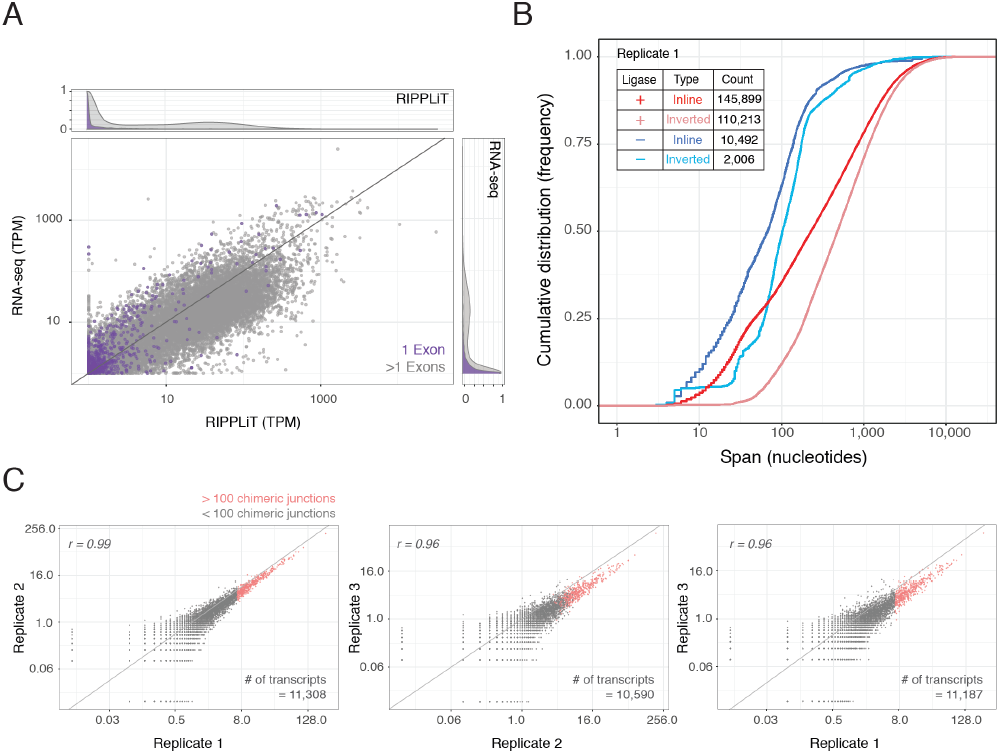
RIPPLiT captures intramolecular ligations in EJC-associated RNAs with high reproducibility. (A) Comparison of fragment coverage (transcripts per million: TPM) for genes with one (purple) or more than one (grey) exon in RIPPLiT (replicate 1 ligase) and RNA-seq (Ge et al., 2016) libraries. Density plots (scaled to 1; top and right margins) show that single exon mRNAs are preferentially depleted from the RIPPLiT library. (B) Cumulative frequency distributions for Replicate 1 of inline and inverted chimeric junction spans + and - ligase. Inset table: Raw junction counts. (C) Scatterplots comparing normalized intramolecular chimeric junction counts per transcript in + ligase libraries among biological replicates. Diagonal line: x=y. Deviations from x=y show that ligation efficiency in Replicate 1 > Replicate 2 > Replicate 3. Red dots, set of transcripts used for scaling plots in Figure 6.

For individual transcripts, the + ligase libraries were strongly enriched for intramolecular chimeric junctions compared to - ligase libraries (**Figure 2C, S2A; Table S2A**). Further, despite differences in ligation efficiency between biological replicates (with + ligase chimeric junction counts in replicate 1 > replicate 2 > replicate 3), the number of chimeric junctions per transcript was highly correlated (r >0.96) across all three replicates (**Figure 2D**). Thus, intramolecular mRNA ligations were abundantly represented in our datasets and exhibited high biological reproducibility.

### RIPPLiT faithfully captures intra-and inter-RNA 3D structure of rRNAs

Due to the high cellular abundance of ribosomes, ~20% of our RIPPLiT fragments mapped to rRNAs, with 18S rRNA exhibiting the highest fragment density (**Table S2B**). Because the folded structure of 18S rRNA is well known, we could use these data to test the validity of our method. Whereas + and - ligase libraries exhibited similar fragment coverage across the entire 18S rRNA (**Figure 3A, tracks along sides of heatmap**), chimeric junctions were 18- to 26-fold more abundant in + ligase than in - ligase libraries (**inset numbers in Figure 3A, B, Table S2B**). This difference is readily apparent in the + and - ligase chimeric junction heatmaps (**Figure 3A**). Further, both inline and inverted chimeric junctions (**Figure 3B**) occurred with nearly equal frequency in the + ligase libraries in a pattern that was reproducible across all three biological replicates (**Figure 3A**).

**Figure 3.**
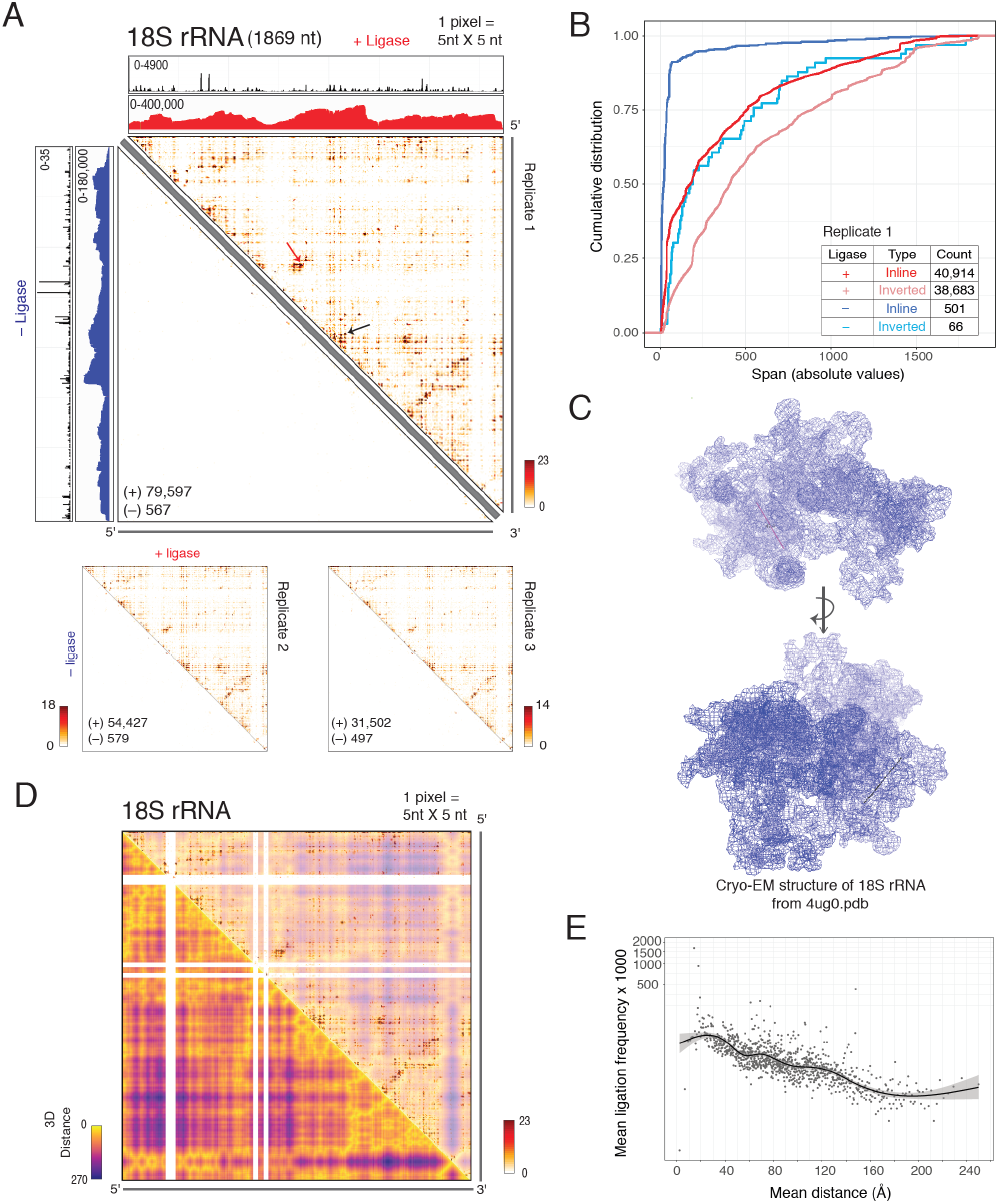
Ligations in 18S RNA occur between 3D-proximal regions. (A) Chimeric junction heatmaps for human 18S rRNA - ligase (lower left) and + ligase (upper right). Pixel color scales, number of junctions per 5×5 nt bin; numbers in lower left corners, number of chimeric junctions obtained per indicated library. Replicate 1 coverage tracks show fragment distributions (red and blue) across the entire transcript and chimeric junction frequencies (black) at each individual nucleotide. Note large scale differences for - and + ligase chimeric junction tracks. (B) Cumulative frequency distributions of inline and inverted chimeric junction spans on 18S rRNA for - and + ligase libraries (Replicate 1). Inset table: Number of inline and inverted chimeric junctions for each library. (C) Structure of 18S rRNA (4UG0 (Khatter et al., 2015)) showing the two chimeric junctions marked with red and black arrows in (A) mapped onto the structure as lines. (D) Heatmaps for mean Euclidean phosphate-phosphate distances (bottom) overlaid with chimeric junctions (top) binned at 5×5 nt. White pixels: regions absence from structure. (E) Scatterplot showing mean ligation frequency in Replicate 1 as a function of mean Euclidian distance for 1,000 bins each containing 79-80 chimeric junctions. Black line shows smoothing (GAM: generalized additive model) with grey area displaying confidence interval (0.95) around smoothing.

Since all rRNA molecules adopt the same 3D structure, 18S chimeric junctions should be most prevalent at locations that are both accessible to nuclease digestion and close enough in Euclidean (3D) space for subsequent ligation. To assess whether our observed junction pattern was consistent with native 18S rRNA folding within the ribosome, we use a high-resolution human 80S ribosome cryo-EM structure (**Figure 3C**; (Khatter et al., 2015)) to calculate mean Euclidean phosphate-phosphate distances for every 5 nt bin. Overlaying the + ligase chimeric junction frequency and mean 3D distance heatmaps (**Figure 3D**) revealed preferential association of chimeric junctions with 3D-proximal regions (yellow) compared to 3D-distal regions (purple). Because ligations require both accessibility and proximity, not all 3D-proximal regions were enriched for chimeric junctions. Nonetheless, a plot of mean 3D distance vs chimeric junction frequency shows the expected decay in junction frequency as Euclidean distance increases **(Figure 3E)**. This same relationship was observed for chimeric junctions mapping within 28S rRNA **(Figure S3)**.

In addition to intramolecular chimeric junctions, we also observed thousands of junctions for which the individual fragments mapped to different rRNA species (**Table S2A**). These intermolecular junctions were 81 to 114-fold more prevalent in + ligase than - ligase libraries (**Table S2B)**, suggesting that most of the + ligase junctions represented true intermolecular ligation events. Mapping these onto the 80S ribosome structure resulted in a similar Euclidean distance versus junction frequency distribution as the intramolecular junctions (**Figure S4A**). In addition, most intermolecular junctions between the 5.8S and 28S rRNA occur near the 5′ end of the 28S rRNA (**Figure S4B**), consistent with the known 5.8S-28S interaction domain (Walker and Pace, 1983) (**Figure S4C**). Thus, RIPPLiT accurately reports 3D structural information in native RNPs, and can capture both intra- and intermolecular interactions within stable multi RNA complexes with a high signal to noise ratio.

### No evidence for stable intermolecular interactions between Pol II transcripts

More than 80% of the >10,000 spliced Pol II transcripts represented in our datasets had more than one unique intramolecular chimeric junction in the + ligase libraries (**Figure 4A, Table S3**). Further, the number of unique intramolecular chimeric junctions per transcript was substantially higher when ligase was included (compare the purple and green distributions). In contrast, intermolecular junctions were rare; only 5% of transcript pairs exhibiting apparent intermolecular junctions had more than one unique junction, and the cumulative histograms were nearly identical between the + and ligase libraries (compare the blue and pink distributions). This suggests that the vast majority of intermolecular junctions on Pol II transcripts were due to mapping artifacts. Thus, we found no compelling evidence for biochemically stable intermolecular interactions between different mRNA species (**Figure S2B, C**; **Figure S5**).

**Figure 4.**
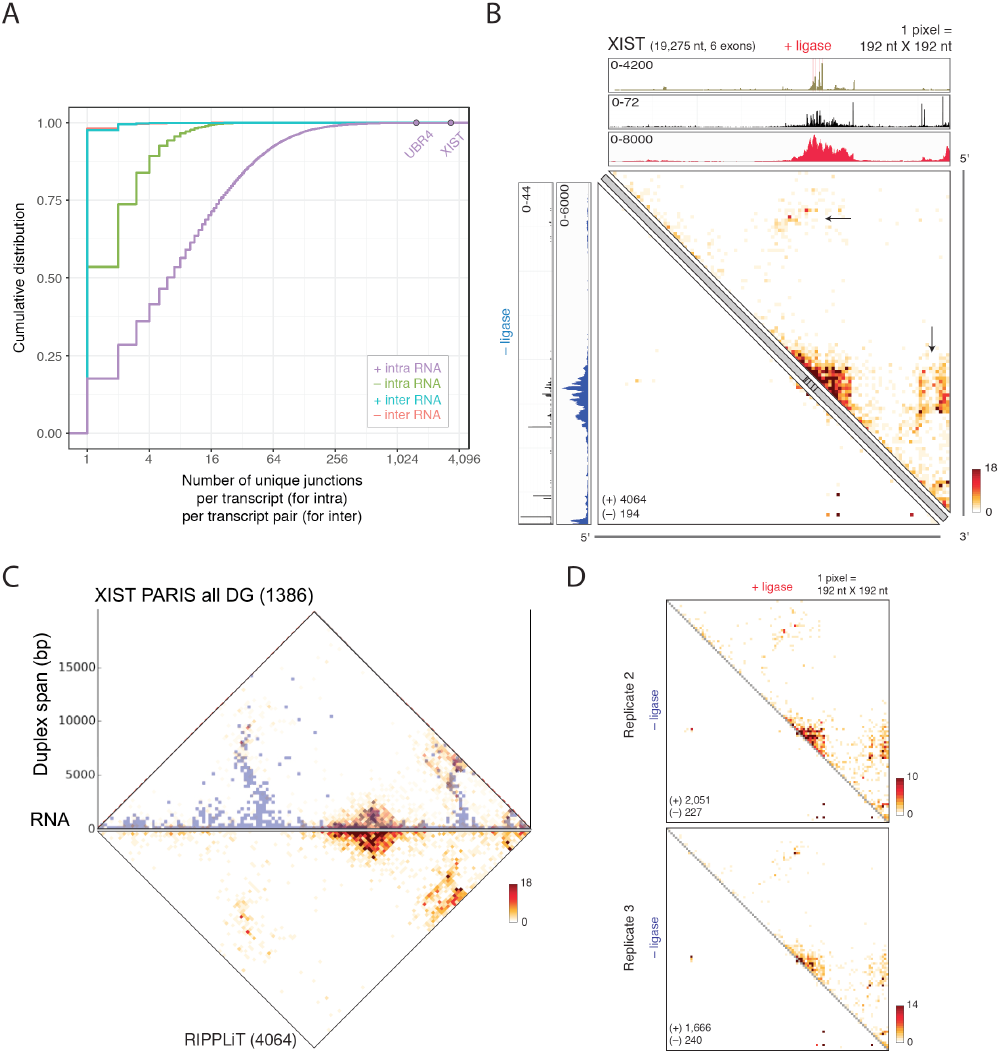
RIPPLiT captures higher-order structure of ncRNA-XIST. (A) Cumulative distribution of unique intramolecular and intermolecular chimeric junctions per RNA Pol II transcript. XIST and UBR4 marked for reference. (B) Chimeric junction heatmaps for XIST (Replicate 1). All is as in Fig. 3A except that tick marks along diagonal gray bars indicate exon-exon junctions. Topmost coverage track displays EJC long footprint coverage obtained from EJC RIPiT experiments (Singh et al., 2012), with pink lines indicating exon-exon junctions. Black arrows indicate long range interactions within the first and the last exon. (C) PARIS (Lu et al., 2016) Duplex Groups (DGs) overlaid with RIPPLiT data for XIST. Numbers inside brackets indicate the number of junctions obtained in each dataset. (D) XIST chimeric junction heatmaps for Replicates 2 and 3.

### RIPPLiT captures XIST structure

The most highly represented Pol II transcript in our datasets was XIST, a long non-coding RNA that functions in X chromosome inactivation. Consistent with its high abundance and nuclear retention (meaning that no EJCs are stripped away by cytoplasmic translation), XIST had ~2.5-fold more chimeric junctions than any other Pol II transcript. XIST consists of two unusually long terminal exons (~7 kb and ~11 kb) flanking four smaller (64 to 209 nt each) internal exons. Reflecting this primary structure, mapped fragments in both - and + ligase libraries exhibited the greatest depth on and around the four internal exons (**Figure 4A**). Chimeric junctions were ~7.5-fold more abundant in + than - ligase libraries (**Table S4**), with + ligase junctions indicating the existence of short- (spanning <200 nt; typically occurring within one exon or between immediately adjacent exons), mid (spanning 200-500 nt) and long-(spanning >500 nt) range interactions. Prominent long-range interactions occurred between the internal exons and the region immediately adjacent to the polyadenylation site in exon 6, and between the internal exons and a region close the 5′ end of exon 1 (**Figure 4A; black arrows**). This pattern was highly reproducible across all three biological replicates (**Figure 4D**). PARIS, which uses in-cell psoralen crosslinking to identify base paired regions (Lu et al., 2016), has indicated the presence of two large secondary hairpin structures in XIST exons 1 and 6 (**Figure 4B**). Consistent with the existence of these structures, RIPPLiT captured interactions at the bases of the hairpins between regions otherwise distant in nucleotide space. Another RNA secondary structure probing method, SHAPE-MaP (Smola et al., 2016), measures 2¢-OH accessibility and flexibility in cells. Compared to purified, protein-free XIST RNA folded *in vitro*, the region around the internal exons is substantially less reactive to the SHAPE reagent in cells (Lu et al., 2016; Smola et al., 2016). Because little secondary structure exists in this region, decreased SHAPE reactivity in cells is best explained by strong intracellular RNA-protein interactions, such as EJCs and associated factors. In this same region, RIPPLiT yielded numerous short to medium range chimeric junctions, suggestive of substantial RNA compaction. Combined, these datasets illuminate the higher order structure of XIST RNA in cells. Whereas PARIS reveals secondary structural elements and SHAPE-MaP reveals potential protein interaction domains, RIPPLiT reveals long-range interactions between these features, including interactions between the internal exons and sequences flanking the hairpins. RIPPLiT thus provides a powerful new addition to the toolkit for transcriptome-wide 3D structure probing of native RNPs.

### Chimeric junctions evenly distribute across mRNAs

We next focused on the set of 490 mRNAs exhibiting ≥100 intramolecular ligation events in at least one biological replicate (**Figure 2C**, **Table S5**). In contrast to XIST, individual fragments in both + and −ligase libraries generally distributed all across the entire length of spliced mRNAs. Exemplifying this were UBR4 (~16 kb; 106 exons), TRRAP (~12 kb; 70 exons), SPEN (~12 kb; 15 exons); PRPF8 (~7 kb; 43 exons), and EEF2 (~3 kb; 15 exons) (**Figure 5A-C and Figure S6**). The pattern of chimeric junctions was similarly distributed across the entire length of each mRNA, indicating a general absence of locus-specific structures. In rRNAs and XIST, such locus-specific structures result in strong chimeric junction enrichments between specific regions (**arrows in Figure 3A and 4A**). In contrast, chimeric junctions on mRNAs were homogeneously distributed across the entire interaction map, with only a general inverse relationship between chimeric junction formation and nucleotide distance. This dispersed pattern was observed in all three biological replicates (**Figure S7A**). Further, this pattern was not simply attributable to read density differences, as XIST + ligase replicate 3 and UBR4 + ligase replicate 1 have a similar number of chimeric junctions, but very different interaction maps (**Figure S7B**). Notably, we did not observe elevated chimeric junctions spanning entire lengths of mRNAs, which would be expected if the 5’ and 3’ ends were closely juxtaposed in a circular arrangement.

**Figure 5.**
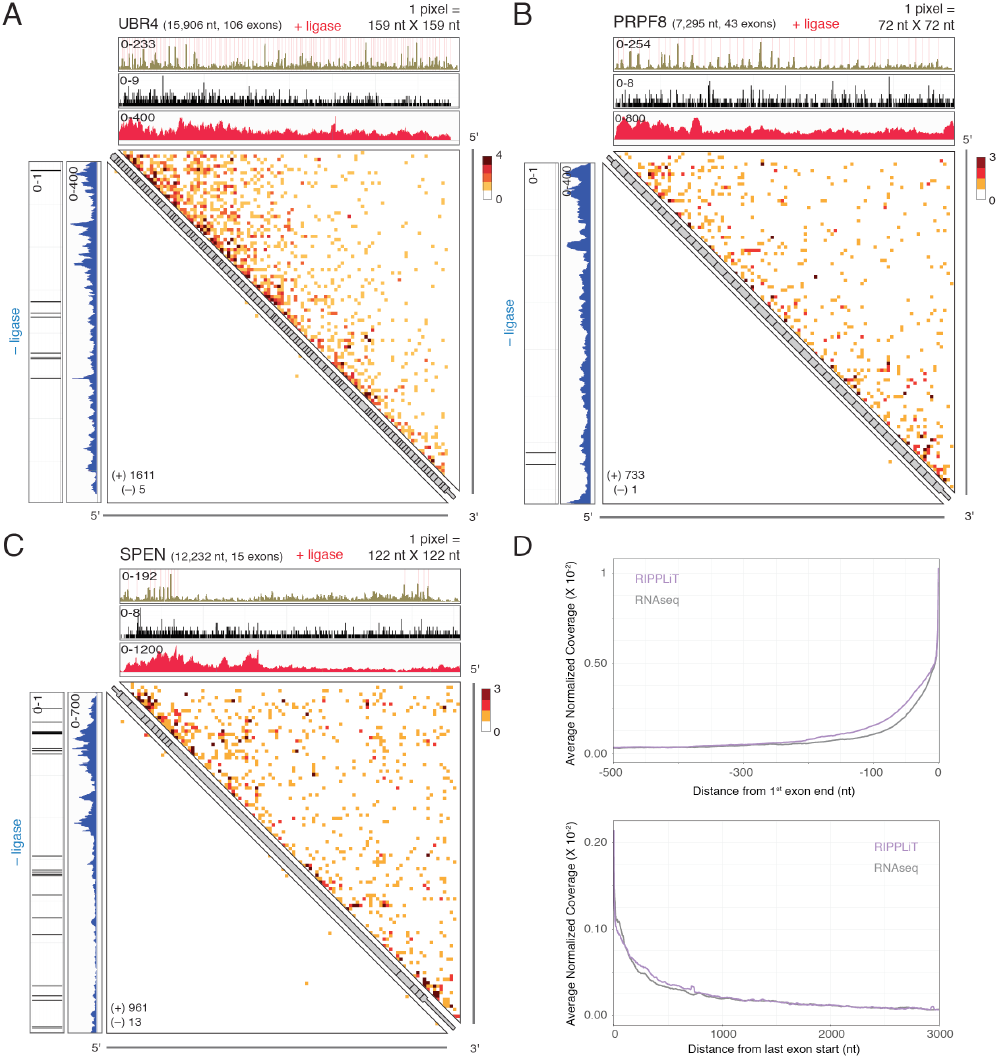
RIPPLiT captures higher-order structure of spliced Pol II transcripts. (A-C) Chimeric junction heatmaps for spliced mRNPs. All is as in Fig. 2A except that tick marks along diagonal gray bars indicate exon-exon junctions. In A-C, coding exons and UTRs are thicker and thinner sections, respectively. Topmost coverage track (tan) displays EJC long footprint coverage obtained from EJC RIPiT experiments (Singh et al., 2012), with pink lines indicating exon-exon junctions. (D) Aggregate normalized coverage plot for RIPPLiT Replicate 1 (purple) and RNA-seq (Ge et al., 2016) (grey) libraries across first exon and last exons for the set of transcripts with >100 chimeric junctions in at least one biological replicate (**Figure 2C**).

Another notable feature was the coverage across an exceptionally large internal exon in SPEN (**Figure 5C**). Whereas the other internal SPEN exons ranged from 101 to 483 nt, exon 11 spans 8,176 nt. Nonetheless, mapped fragments and chimeric junctions were observed throughout this entire exon, with read density on this exon being ~12-fold greater than read densities on XIST exon 1, a segment of similar length. Thus, the structural scaffold of which the EJC is a part appears to encompass all internal mRNA exons regardless of length. Comparable decays in RIPPLiT and RNA-seq coverage on first and last exons indicate that this scaffold also extends into the the terminal exons (**Figure 5D**).

### Polymer analysis indicates that pre-translational mRNPs form flexible rods

The relatively even distribution of chimeric junctions across the interaction map indicates a general absence of strong locus-specific secondary structures within nascent mRNPs. Rather the mRNAs behave more like flexible polymers. This is consistent with in-cell SHAPE (Spitale et al., 2015) and DMS-seq (Rouskin et al., 2014) data, which indicate that mRNAs in cells are more flexible (i.e., less structured) than *in vitro*-folded protein-free mRNAs. Such higher flexibility could reflect either loose packing of the mRNA strand or tight packing without specific points of contact. Without specific contact points provided by base pairing interactions, each mRNA molecule will adopt a slightly different mRNP structure in which different nucleotides are brought into close proximity, thus resulting in an even distribution of chimeric junction in the ensemble population.

Another feature of RIPPLiT data on mRNAs is the inverse relationship between ligation frequency and primary sequence span (**Figure 5A-C**). Both the even distribution of interactions across the entire length of mRNAs and the general decay in ligation frequency with increasing nucleotide span are reminiscent of proximity ligation maps for chromatin. As with Hi-C interaction heatmap for DNA (Lieberman-Aiden et al., 2009), polymer analysis of the ensemble data can provide insight into RNA packaging. Quantitative analysis of the relationship between contact frequency and the number of monomer units separating individual contact points can reveal the conformational organization of the polymer (e.g., whether or not it is a random coil (Fudenberg and Mirny, 2012)). We therefore calculated average chimeric junction frequency as a function of nucleotide distance for all 490 mRNAs. To enable data aggregation across different length mRNA species, we divided each transcript into 100 bins (1% per bin) and calculated the fraction of total chimeric junctions in each bin. We could then create composite matrices by first summing the normalized junction frequencies across all bins for multiple mRNA species and then balancing the matrix using Iterative Correction as has been used for analysis of Hi-C data (Imakaev et al., 2012; Rao et al., 2014). Since, the resultant composite matrices represent average ligation frequencies across many transcripts and balancing will mitigate any biological (e.g., the locations of specific EJC and EJC-associated protein binding sites in any one mRNA) or technical (e.g., read mappability) biases.

Since mRNA length varies widely, it seemed possible that mRNPs could adopt different structures at different length scales. To test this, we removed the most extreme length outliers (5% shortest and longest), divided the remaining 442 transcripts into three equal member length groups (short: 2,403-4,683 nt; medium: 4,683-6,862 nt; and long: 6,870-13,492 nt), and created composite heatmaps for each group (**Figure 6A**). Strong similarity between all three heatmaps suggests no major structural differences driven by length. Further, as we had also observed for individual mRNAs (**Figure 5A**), there was no evidence of stable interactions between the 5′ and 3′ ends (as indicated by the absence of signal in the upper right and lower left corners of heatmaps in **Figure 6A**).

**Figure 6.**
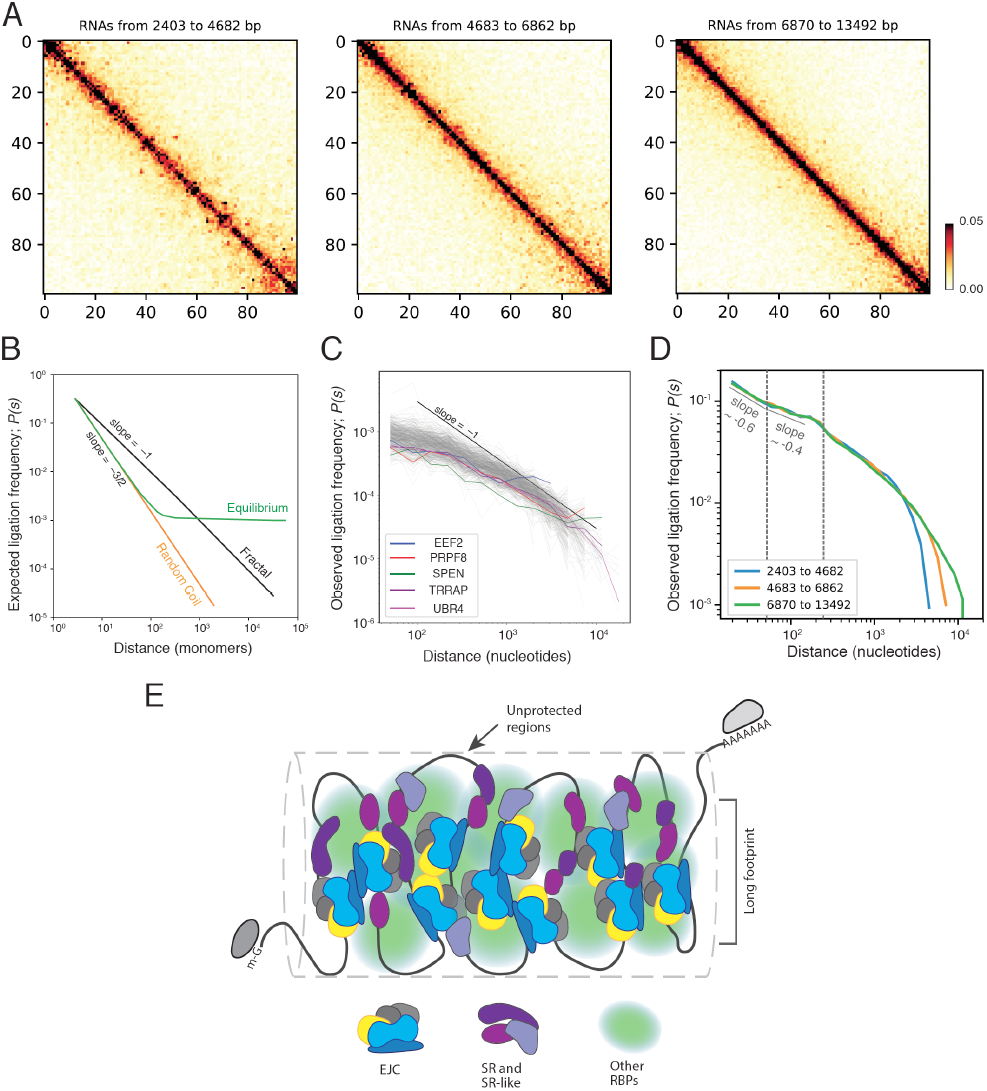
Within mRNPs, mRNAs are densely packed into linearly organized rod-like structures. (A)Metagene heatmaps for transcripts of indicated length ranges. These heatmaps were created by dividing each transcript into 100 bins (1% length per bin = 1 pixel) and normalizing each heatmap to a total chimeric junction frequency of 1.0, thus enabling data integration across multiple mRNA species of differing lengths. Color scale: Normalized mean chimeric junction frequency per pixel. (B)Expected scaling plots for different polymer types. Probability of contact; P(s) (equivalent to ligation frequency) is plotted against distance (here, monomers constituting the polymer) on a log-log axes. EG, equilibrium globule; RC, random coil; FG, fractal globule. (C)Mean observed scaling plot for 490 transcripts (grey lines) with more than 100 chimeric junctions in at least one biological replicate. Data from all replicates were combined and mean ligation frequency plotted as a function of distance in nucleotides on log-log axes. Each color represents a transcript (exemplifying different lengths) shown in Fig. 6A and Fig. S6. Black line indicates a slope of −1. (D)Mean observed scaling plot of all replicates combined for each length group shown in Fig. 6A. Vertical dotted lines indicate approximate boundaries for exponent changes. (E)Model of rod-like mRNP, with EJCs and other mRNP proteins protecting large RNA regions from nuclease digestion.

The exponent (slope) by which the frequency of chimeric junctions decays with nucleotide distance (span) on a log-log scaling plot can reveal properties of the folded state of the RNA polymer (**Figure 6B**; (Fudenberg and Mirny, 2012)). For instance, if mRNAs fold as simple free random coils, as has been proposed based on in cell SHAPE data (Rouskin et al., 2014; Spitale et al., 2015), one would expect an exponent of −3/2, independent of mRNA length. Conversely, if mRNAs fully equilibrate to form equilibrium globules, proximity ligation frequency should initially decay with an exponent of −3/2, but then reach a plateau where ligation frequencies become independent of nucleotide distance. Neither feature was observable in our data, either for individual mRNAs (**Figure 6C**) or for composite of the three length groups (**Figure 6D**). Thus, the paths assumed by mRNAs in nascent mRNPs are neither simple random walks nor equilibrium globules.

Three main features of the composite mRNA scaling plots are: (1) an initial slow decay (exponent ~-0.6) in ligation frequency for spans up to 40 nucleotides; (2) an even slower decay (exponent ~-0.4) for spans of 40-250 nucleotides; and then (3) a steady increase in the exponent as spans increase beyond 250 nucleotides. These three features are remarkably similar to the Hi-C interaction patterns observed for prophase chromosomes lacking condensin **II** (see Figure 5D in (Gibcus et al., 2018)), albeit on very different length scales. Such chromosomes are predicted to form highly packed and elongated structures containing numerous short (60-80 kb) loops connected by condensin I. The initial slow decay phase in the Hi-C scaling plots reflects interactions within individual loops, the second phase reflects contacts between adjacent loops, and the third phase (where the decay increases with span) is indicative of an elongated rod-like structure wherein interloop interactions become increasingly rare at very long distances.

For mRNAs, all three length classes exhibited identical first and second phases, with the very small exponents indicating a highly packed RNA conformation. The third phase was equally indistinguishable up to the point where the drop off sharply increased as spans began to exceed mRNA length in each class. Thus, consistent with the heatmaps, we observed no evidence of specific 5′ and 3′ end interactions in any length class. Rather the increasingly steep decay at larger nucleotide distances is indicative of a flexible elongated structure. We therefore conclude that spliced pre-translational mammalian mRNAs are compacted linearly into densely packed non-circular rod-like structures irrespective of mRNA length (**Figure 6E**).

## Discussion

Here we describe a new proximity ligation technique, RIPPLiT, to capture higher order structural information in RNPs of defined protein composition. By enriching for EJC-associated RNAs, we were able to investigate the overall architecture of spliced Pol II transcripts in association with stably-bound proteins acquired during transcription and pre-mRNA processing. Our data indicate that prior to translation, spliced mRNAs and their associated proteins are tightly packed into linearly-organized flexible rod-like structures irrespective of length. Within these structures, we could detect no specific interaction between the 5¢ and 3¢ ends.

### RIPPLiT captures 3D RNP structural information independent of base pairing

Proximity ligation coupled with semi-quantitative PCR has been used to map chromatin interactions and to delineate the 3D conformation of yeast chromosomes (Dekker et al., 2002). Subsequent technical developments allowing identification by proximity ligation of chromatin interactions genome-wide enable interrogation of the overall 3D arrangement of DNA in cells, and such studies are starting to reveal the folding principles of chromatin in different biological states at high resolution (Gibcus et al., 2018; Lieberman-Aiden et al., 2009). The first adaptation of proximity ligation to RNA was CLASH (Cross-linking, Ligation, And Sequencing of Hybrids; (Kudla et al., 2011)), which captured intermolecular RNA-RNA base pairing interactions (e.g., miRNA-mRNA) associated with a particular UV-crosslinked RBP (e.g., Ago1) (Helwak et al., 2013). Multiple subsequent studies have described methods for capturing intra-and intermolecular base-pairing interactions transcriptome-wide, either by psoralen crosslinking (e.g., PARIS (Lu et al., 2016), SPLASH (Aw et al., 2016) and LIGR-Seq (Sharma et al., 2016)) or by UV-crosslinking to the double-stranded RBP, Staufen 1 (HiCLIP; (Sugimoto et al., 2015)). Base-pairing interactions also dominate datasets obtained from other proximity ligation methods (RPL (Ramani et al., 2015) and MARIO (Nguyen et al., 2016)) even though these methods were not specifically designed to capture secondary structures. Therefore, previous proximity ligation methods for probing higher order structures within RNPs mainly captured interactions driven by base pairing, and so were highly skewed toward ncRNAs. In contrast, we enriched for EJC cores, deposition of which requires single-stranded RNA (Andersen et al., 2006; Mishler et al., 2008; Singh et al., 2012). Although our data do include some interactions driven by base-pairing (e.g., those at the base of large secondary structures in XIST; arrows in **Figure 4B**), more common were interactions in regions without strong base-pairing tendencies. Thus, a major difference between previous proximity ligation methods and RIPPLiT is that we capture higher order interactions in RNPs formed as a consequence RBP binding and packaging as well as base-pairing. This coupled with the high coverage throughout mRNAs (**Figure 5**) allows for interrogation of the overall 3D organization of stable RNA-protein complexes.

The strong enrichment of our EJC RIPPLiT datasets for spliced Pol II transcripts and the high depth of chimeric junctions in the + ligase libraries (464,426 unique junctions on Pol II transcripts in all) allowed us to investigate the 3D structural properties of hundreds of spliced mRNAs. Notably, the chimeric junctions we observed on mRNAs were almost entirely intramolecular (**Figure 4A**). Despite intensive examination of our data, we found no convincing evidence for any specific intermolecular mRNA-mRNA interactions (**Figures S2 and S5**). Even though intermolecular interactions between rRNAs were readily apparent in our datasets (**Figure S4**), we could detect no specific multi-RNA mRNPs. Nonetheless, intermolecular interactions between different mRNA species have been reported in previous proximity ligation studies that employed in situ crosslinking in yeast, HeLa cells, lymphoblastoid cells, hESC line H1 (Aw et al., 2016) and mouse embryonic stem (ES) cells (Nguyen et al., 2016), albeit at extremely low counts per interaction. Thus, if intermolecular mRNA interactions do occur in HEK293 cells, they are either exceedingly rare or they cannot survive the affinity purification and T1 RNase digestion conditions employed here. Of note, there is evidence for the existence of large, multi-mRNA transport granules in cells with highly extended processes (e.g. neurons and oligodendrocytes) (Carson et al., 2008; Park et al., 2014). It will therefore be of interest to apply RIPPLiT or another proximity ligation approach to these cell types both to investigate the mRNA makeup of these granules and to determine whether they are pre-or post-translational complexes.

Finally, whereas the majority of chimeric reads in our datasets were indicative of a single ligation event (i.e., two fragments ligated together), numerous reads contained three or more ligated fragments coming from the same mRNA species, and thus likely from a single mRNA molecule. Such reads could potentially provide valuable information as to the overall structure and flexibility of individual mRNP complexes.

### Structural features of mRNPs revealed by RIPPLiT

Our previous study of EJC footprints on spliced Pol II transcripts focused primarily on short (12 to 25 nt) footprints expected for monomeric EJCs (Singh et al.,2012). In that study, we subjected EJC-bound RNAs to high concentrations of RNase I. Yet despite this stringent digestion, the predominant fragment sizes obtained were 30-150 nt, much longer than the EJC or any other known RBP footprint. This indicated that, when associated with their complement of stably-bound proteins, spliced Pol II transcripts are highly protected from RNase digestion.

In the current study we subjected EJC-containing RNPs to milder RNase T1 digestion, both with the intent of preserving overall RNP structure and to generate single-stranded RNA ends with sufficient toeholds for T4 RNA ligase I. For spliced mRNAs, we observed fragment coverage over almost the entire transcript (**Figures 5 and S6**). This is consistent with previous mapping data indicating canonical EJC binding upstream of exon junctions plus strong association with other non-canonical sites, most likely through protein-protein interactions with other RNA-binding proteins (Sauliere et al., 2012; Singh et al., 2012). This relatively homogenous fragment coverage, even across extremely long internal exons (**Figure 5C**), indicates that canonical EJCs are just one small part of a much larger, highly stable multicomponent interaction network.

Scaling plots (**Figure 6**) strongly suggest that mammalian pre-translational mRNAs exist as linearly organized rod-like structures. Within these rods, EJCs and associated proteins likely form a stable scaffolding that nucleates and maintains the mRNA in a densely packaged state. Our current data combined with previous imaging studies and our findings that mRNPs are strongly protected from RNase digestion suggest a very compact structure with many different species contributing to a dense network of protein-RNA and protein-protein interactions. The high frequency of chimeric junctions with spans less than 40 nt likely represents ligations occurring between the ends of short excursions from this tight proteinaceous core.

Rod-like packaging, which likely occurs in a first-come-first-served order during mRNA synthesis and processing, has multiple biophysical and mechanical advantages. First, by preventing RNA knots and by compressing exons into a form less prone to physical breakage than extended RNA, compacted rods can preserve the functionality and integrity of newly synthesized messages. Second, rod-like nanoparticles diffuse more rapidly than spherical particles through adhesive polymeric gels, such as provided by the dense polymeric environment of the nucleus with its innumerable weak interaction sites for RNA-protein complexes (Wang et al., 2018). Third, rods of uniform thickness rather than spheres of varying diameter dependent on mRNA length have clear advantages for passage through the nuclear pore complex. Intriguingly, a recent intracellular tracking study following differently shaped nanoparticles reported that even when equal diameter objects are compared, rod-and worm-shaped nanoparticles traverse nulear pores more efficiently than spheres (Hinde et al., 2017). Consistent with this, even globular Balbiani ring mRNPs can be seen passing through nuclear pore as rods (Skoglund et al., 1983).

Finally, mRNAs undergoing translation often have functional interactions between the 5′ and 3′ ends (i.e., between proteins bound to the cap and polyA tail) (Christensen et al., 1987; Wells et al., 1998). However, we observed no evidence for such interactions in pre-translational mRNPs. That the ends of mammalian mRNPs do not interact prior to translation is supported by a recent in cell single molecule FISH study, which found no colocalization in the nucleus of probes hybridizing to 5′ and 3′ ends of three different mRNAs (Adivarahan et al., 2017). Interestingly, in that study, 5′-3′ end interactions were observable in the cytoplasm only after polysome collapse upon puromycin treatment. Thus, functional circularization of mRNAs may only occur within polysomes, not in pre-translational mRNPs.

## Acknowledgments

We thank members of the Moore and Dekker labs for discussions and critiques and Daniel Zenklusen for communicating results prior to publication. This work was supported by funding from HHMI, NIH RO1-GM53007 (M.J.M.), and NIH HG003143 (J.D). J.D. is an HHMI Investigator and M.J.M. was an HHMI Investigator during the time this work was conducted.

## Author Contributions

M.M. and M.J.M. originally conceived the project, with M.M. executing all wet bench experiments. B.R.L. and H.O. conceived and wrote ChimeraTie and M.M. implemented all computational analyses with input from M.J.M and J.D. Polymer analysis was done by M.I. under the guidance of J.D. and L.A.M. All authors contributed to data analysis and interpretation. With input from all authors, M.M., J.D. and M.J.M. were primarily responsible for writing the paper.

